# Effects of feeding water hyacinth (*Eichhornia crassipes*) fodder with or without commercial concentrate on zoo-technical performance and profitability in tropical goats and sheep

**DOI:** 10.1101/2024.10.08.617183

**Authors:** Yared Fanta, Yisehak Kechero, Nebiyu Yemane

## Abstract

The utilization of unconventional feed resources, such as water hyacinth is an effective strategy to address feed shortages in tropical and subtropical regions, particularly where access to conventional feeds is limited. In this study, tropical sheep (Doyogena rams) and goats (Woyito-Guji bucks) fed diets containing different amounts of WH were examined for their zoo-technical performance and profitability sing a 2 × 4 × 4 randomized crossover design with two animal species, four nutritional treatments, and four feeding intervals. The dietary treatments consisted of 50% hay + 0% WH + 50% commercial concentrate (CC, T1), 50% hay + 12.5% WH + 37.5% CC (T2), 50% hay + 25% WH + 25% CC (T3), and 50% hay + 37.5% WH + 12.5% CC (T4). The findings showed that compared to goats, sheep had the highest energy and nutrient intake (P<0.001), nutrient digestibility (P<0.001), average daily gain (ADG, g/day), and body weight change (BWC (kg) (P<0.05). Regarding energy and nutritional intake, there was a substantial difference (P < 0.001) between treatment groups for both animal species, with the exception of goats’ consumption of DM, OM, CHO, GE, and ADL (P <0.05). Likewise, significant differences existed between treatment groups for nutritional digestibility, ADG, BWC, and FCE for both species (P <0.001). Moreover, significant interactions (P < 0.005) were seen in all energy and nutrient intake parameters between species and treatment. Furthermore, in tropical sheep and goat breeds, water hyacinth can replace up to 37.5% of the commercial concentrate used for growth and fattening, but it has a major comparative effect on sheep. Feed prices for the T4 group fed sheep and goats were 37.2% and 36.8% lower, respectively, than for the T1 group. Therefore, farmers in the tropics who cannot afford commercial concentrates can still benefit economically by using the dry biomass of water hyacinth in their diet, either with or without it.

## 1. Introduction

Livestock production is a cornerstone of the Ethiopian economy, significantly contributing to food security, livelihoods, and cultural practices (Amejo, 2024). However, the sector constrained by both technical and institutional factors, of these challenges feed scarcity, seasonal fluctuations of feed supply, and increasing competition for traditional grazing lands takes the lion share. These issues are especially pronounced during dry seasons, negatively impacting livestock health and overall productivity, thereby threatening the livelihoods of those who depend on this vital resource (Tonamo *et al*., 2015).

There is an urgent need for creative ways to improve animal nutrition and overall output as the population increases and climate change intensifies. In this situation, it is more important than ever to investigate non-traditional feed sources, especially in rural areas where access to conventional feed is still scarce. One such unconventional feed source is water hyacinth (*Eichhornia crassipes*), an aquatic plant that has garnered attention for its potential as a high-quality feed resource (Hossain *et al*., 2015). Originating from South America, water hyacinth has spread too many parts of the world, including Ethiopia, where it often grows in abundance in lakes, rivers, and wetlands (Stroud, 1994). This invasive species, while posing environmental challenges, offers nutritional benefits that can be harnessed for livestock feeding. Water hyacinth is rich in protein, fiber, and essential nutrients, making it a viable alternative to traditional feed ingredients such as hay and grass (Farid et al., 2016; Suleiman *et al*., 2020; Fanta *et al*., 2024).

Different animal species exhibit varying capacities for digesting and utilizing different feed types, largely influenced by their natural feeding behaviors (Ketshabile, 2008). This is particularly relevant in the context of the browser/grazer dichotomy, which distinguishes between animals that primarily consume leaves and shrubs (browsers) and those that prefer grasses (grazers). Browsers, such as Woyito-Guji bucks, are adapted to extracting nutrients from woody plants, while grazers, like Doyo gena rams, are optimized for consuming herbaceous grasses. This inherent difference in feeding behavior can significantly affect how efficiently these animals utilize alternative feed resources like water hyacinth.

This study aims to compare the zoo technical performance and economic viability of Woyito-Guji bucks and Doyo gena rams of Ethiopian origin/tropical goat types/ when fed diets containing water hyacinth. By examining how the unique dietary preferences and physiological adaptations of these two breeds influence their ability to digest and metabolize water hyacinth, this research will provide valuable insights into their potential roles in sustainable livestock management. The findings will contribute to a deeper understanding of how different livestock species can effectively incorporate unconventional feed sources into their diets, thereby promoting more sustainable feeding practices. Moreover, the significance of this research extends beyond academic interest; it holds practical implications for smallholder farmers in the tropics. Farmers can choose feed options that improve animal health, production, and profitability by learning the best way to use water hyacinth for small ruminants in the area. Additionally, incorporating non-traditional feeds like water hyacinth into livestock diets can assist farmers relieve some of the financial strain they are facing in a time when conventional feed resources are growing more expensive and scarce.

## 2. Materials and Methods

### 2.1. The Experimental Area

The experiment was conducted at Arba Minch University livestock research Farm, southern Ethiopia, from October 2021 and February 2022. Geographically, the study area is located at the base of the western side of the Great Rift Valley in Ethiopia, at 6°2’21”N latitude and 37°34’24”E longitude, and 435 km south of Addis Ababa, Ethiopia. The altitude of the experimental site is 1285 m above sea level. The average annual rainfall varies between 598 mm and 1245 mm, while temperatures range from 15.8 °C to 33.5 °C (Ayda *et al.,* 2021.

#### 2.1.1. Abundance of water hyacinth in Lake Abaya

Water hyacinth (*Eichhornia crassipes*) is a perennial aquatic weed, known for its rapid growth and ability to form dense mats, which present a significant challenge to water bodies in tropical and subtropical regions. Impacting rivers, dams, lakes, and irrigation channels in Lake Abaya alone, this invasive species covers staggering 3500-3700 hectares studies for 2020 (Alemante et al., 2021). In Lake Abaya, Ethiopia, water hyacinth’s invasion has led to a decline in native macrophyte species, while promoting the dominance of water hyacinth itself, impacting the overall ecosystem (Mengistu et al., 2017). The sheer volume of water hyacinth biomass produced annually is remarkable, reaching 150 to 200 tons per hectare (Dymond, 2002), which was equivalent to up to 740,000 tons per entire lake area per year.

### 2.2. Animal care

The experiment was approved by the animal research ethics review committee of Arba Minch University (AMU/AREC/4/2016) following guidelines of the European Union directive number 2010/63/EU (2010) regarding the care and use of animals for experimental and scientific purposes.

### 2.3 Experimental animal and management

The feeding trial was conducted using 24 intact yearling lambs and bucks (12 Doyogena lambs and 12 Woito-Guju bucks) with average weights of 20.78±0.4kg and 19.23±0.3kg, respectively, which were purchased from the local market. The birth histories of animals were known by an interview made during the purchase process. The experimental animals were kept for four weeks in isolation pen and acclimation prior to the commencement of the experiment (Kechero *et al*., 2016), when they were sprayed with an accaricide (diazzinole), dewormed with a broad-spectrum anthelmintic (albendazole) and vaccinated against anthrax and pasteurelosis. During the acclimatization phase, the animals were allowed to adjust to their new environment and were gradually introduced to the experimental diets to facilitate adaptation of their rumen microbes. Each animal was ear-tagged and assigned to individual metabolic pens. Experimental animals were randomly allocated to treatment groups, with feed provided in equal portions twice daily, along with access to clean drinking water and salt licks *ad libitum*. Throughout this period, the animals were weighed after an overnight fast to avoid gut content variation (Kechero *et al*.,2016). Diet allowance for the next period was recalculated according to body weight (BW), and the average weight was recorded as the initial body weight.

### 2.4. Experimental design and dietary treatments

The design of the experiment was randomized a 2×4×4 crossover design with two animal species (sheep & goats), 4 dietary treatment, and 4 feeding periods comprising three lambs and three bucks as a replicate in each period. Experimental animals were given a total of 124 days of feeding regime for the experimental diets, after which 21 days of adaptation for digestion trial, and to remove the carryover effects and 10 days of feces collection for 4 sequential periods. The dietary treatment allocations for the subsequent period were adjusted based on the animals’ live body weight. The daily experimental diet (comprising both basal feed and supplements) was administered to experimental animals that was set at 50 g dry matter/kg live body weight, ensuring a minimum of 8.36 MJ/kg of metabolizable energy and 70 g CP/kg on a dry matter basis (NRC, 2007). Water hyacinth was harvested from Lake Abaya in the shoreline of Arab Minch Zuria district of Chano Mile ward. The commercial concentrate was purchased from Muza livestock feed maker and distributor, a private enterprise in Arba Minch Town. The basal diet, mixed plant hay plant came families like Asteraceae, Fabaceae, Poaceae, Commelinaceae, Cypraceae, and others, with Fabacea, Poaceae, and Asteraceae found to be 85% total plant composition) was harvested from the Arba Minch University research farm. There was free access to water for experimental sheep and goats. The water hyacinth and commercial concentrate (Table 1), and their mixtures were offered as dry matter basis. A list of treatment groupings can be seen in Table 1.

**Table 1.** Lists and composition of experimental dietary treatment combinations.

|  | **T1** | **T2** | **T3** | **T4** |
| --- | --- | --- | --- | --- |
| Grass hay | 50% | 50% | 50% | 50% |
| Concentrate | 50% | 37.5% | 25% | 12.5% |
| *E. crassipes* | 0% | 12.5% | 25% | 37.5% |

### 2.5 Chemical analysis

The samples of water hyacinth, commercial concentrate, hay and refusals that were ground to pass 1 mm mesh size analyzed for DM, ash, and N contents according to the AOAC (2019). The crude protein (CP) was determined by multiplying %N by 6.25, and the organic matter (OM) was estimated by subtracting the ash content from 100%. The neutral detergent fiber (NDF) was analysed according to Van Soest et al. (1991), whereas acid detergent fiber (ADF) and acid detergent lignin (ADL) contents were determined according to the procedure of Van Soest and Robertson (1985) at the analytical chemistry laboratory, Arba Minch University, Ethiopia.

### 2.6 Calculations of energy and digestible nutrient values

The gross energy (GE), digestible energy (DE), and metabolizable energy (ME) of treatment diet were calculated using equations from Hvelpund et al. (1995) as follows:

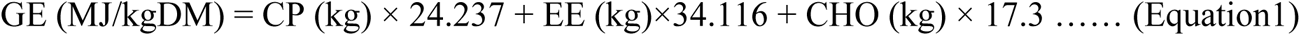

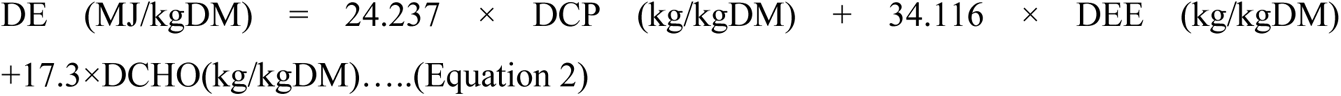

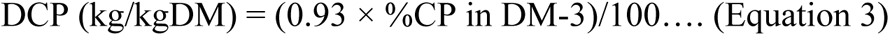

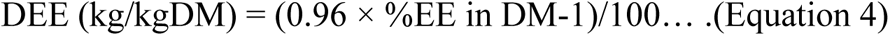

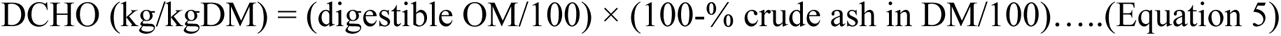

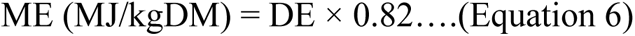

For the estimation of total carbohydrates (total CHO) the following equations were proposed by Sniffen et al. (1992):

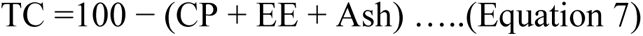

Total digestible nutrient (TDN) of the treatment diet was calculated following (Ranjhnan, 2001).

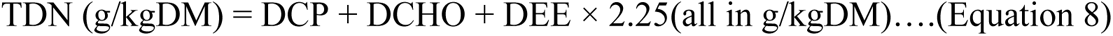

The non-fiber carbohydrate (NFC) was calculated following Jayanegara *et al*. (2019).

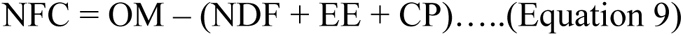

### 2.7. Feed intake

Daily feed intake was calculated by recording the number of feed offers and feed refusals per head (Gemechu et al., 2020). Samples of the feed offered were collected for each batch of feed, and samples of the animal’s refusals were collected per treatment and combined to determine the chemical composition. Daily feed intake of individual animals was calculated as following:

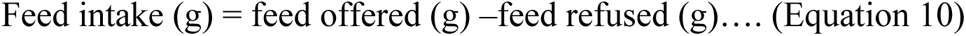

### 2.8. Energy and nutrient digestibility

Prior to the feeding trial, a 10-day digestibility trial had been carried out, promptly followed by a three-day adaptation period to help the experimental animals get used to carrying fecal bags, followed by a complete collection of feces over seven consecutive days. On each morning, before the animals were given feed and water, the entire amount of excrement was gathered by emptying the bags of individual animals. About 20% of the fresh excrement was subsampled of individual animals after it was weighed and thoroughly mixed. It was then kept in a deep freezer at –20 °C. The samples were pooled by animal, and 20% of each pool was subsampled, weighed, and partially dried at 60 °C for 72 hours. Using the following formula (Manaye et al., 2009), the apparent digestibility of energy, and nutrients was calculated.

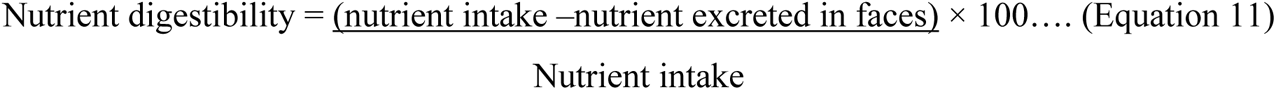

### 2.9. Feed conversion efficiency

Feed conversion efficiency is a measure used to evaluate how efficiently experimental animals transform feed into body weight gain in relation to average daily feed intake. It was measured using the formula suggested by McDonald et al. (2010).

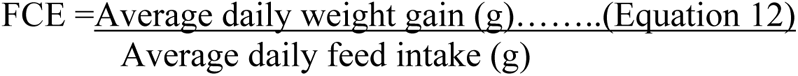

### 2.10. Body weight change and average daily weight gain

Body weights of the animals were measured at the last day of each experimental period, after overnight fasting. Average daily weight (BW) change was calculated on a period basis as the difference between the final and initial BW divided by the number of feeding days. Mean daily body weight change (Kahsu et al., 2021) was calculated as:

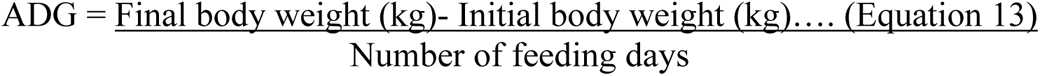

### 2.11. Partial budget analysis

Cost benefit analysis was performed to evaluate the profitability of feeding graded levels of Water hyacinth meal with concentrate mix to Woyto-Guji bucks and Doyo-Gena ram and considering the main cost components. The study used a concentrate mix costing 24 Birr per kilogram, natural grass hay at 10.71 Birr per kilogram, and water hyacinth meal priced at approximately 6.94 Birr per kilogram. Sheep and goat were bought within a price range of 2,000-2,500 birr, and at the end of experiment their prices were determined based on local market values. Nonetheless, the live animals were valued at the beginning and end of the trial by individuals who had previously dealt with sheep and goat trade, and the partial budget analysis included average estimations of the sheep and goats purchase and selling values. The economic viability of different dietary treatments was analyzed using a partial budget analysis, based on the methodology developed by Upton (1979).

It evaluates profit or loss, defining net income (NI) as the difference between total returns (TR) and total variable costs (TVC).

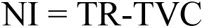

The change in net income (ΔNI) is determined by the difference between the change in total return (ΔTR) and the change in total variable cost (ΔTVC).

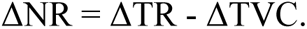

The marginal rate of return (MRR) assesses the increase in net income (ΔNI) that results from each additional unit of cost (ΔTVC).

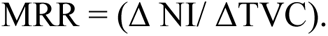

### 2.12. Statistical analysis

Variance analysis was conducted following a 2 × 4 × 4 factorial layout in a randomized cross-over design using mixed model procedures of SAS (2013 version 9.4). Means for different treatments, periods, and animal species variation on zoo technical performances and economic viability of both sheep and goats were separated via the Tukey’s HSD method, and significance differences were declared at P < 0.05. The model applied for all variables indicated below:

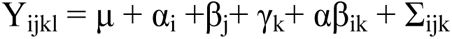

Where, Y_ijkl_ = the response due to the animal i, in period j, treatment k, and interaction effects,

μ = the overall mean effect,

α_i_ = the i^th^ fixed effects of the animal species groups (I = sheep or goat) (subject; i = 1, 2, 3…12),

β_j_ = the j^th^ fixed effects of the experimental period (j = 1, 2, 3, 4),

γ_k_ = the k^th^ fixed effect of the treatment (k =1, 2, 3, 4),

αβ_ik_ = the interaction effect between species i and treatment k, and

Σ_ijkl_ = the random error

## 3. Results

### 3.1. Nutritional quality of experimental feedstuffs

As expected, the *E. crassipes* had 109.07 g DCP/kg DM, had a double fold digestible crude protein content than the hay, which had 50.2 g DCP/kg DM. The hay, primarily composed of fiber, exhibited lower levels of digestible nutrients but contained a high proportion of ADL (72.3 g/kg DM), indicating high fiber content. The commercial concentrate displayed a rich nutrient profile with the highest levels of digestible crude protein (164.2 g/kg DM), total digestible nutrients (419.38 g/kg DM), digestible energy (8.04 MJ/kg DM), and metabolizable energy (6.6 MJ/kg DM), and gross energy (18.7 MJ /kg DM) content of the diets, followed by water hyacinth (*E. crassipes*). With an increase in treatment numbers, there was a tendency for the energy and nutritional values of treatment groups (T1 through T4) to gradually decline in nutrient levels. Treatment (T1), containing the highest proportion of concentrate mix, had the highest DCP (107.2 g/kg DM), TDN (316 g/kg DM), DE (5.95 MJ/kg DM), and ME (4.89 MJ/kg DM) and GE (17.8 MJ/kg DM) values.

**Table 1.** Nutritional value of dietary treatments and experimental diets (g/kg dry matter)

| Energy & Nutrients | Nutritional quality test diets, g /kg |  |  | Nutritional quality of treatments, g/kg |  |  |  |
| --- | --- | --- | --- | --- | --- | --- | --- |
|  | WH | Hay | C.Concentrate | T1 | T2 | T3 | T4 |
| DM | 923.4 | 922.3 | 901.7 | 912 | 915 | 917 | 920 |
| OM | 891.2 | 910.6 | 931.5 | 921 | 916 | 911 | 906 |
| EE | 54.2 | 37.8 | 79 | 58.4 | 56 | 52.2 | 49.1 |
| DCP | 109.1 | 50.2 | 164.2 | 107 | 100 | 93.4 | 86.5 |
| CHO | 71.65 | 81.56 | 67.27 | 74.4 | 75 | 75.5 | 76.06 |
| NFC | 19.27 | 15.27 | 28.8 | 22.1 | 20.8 | 19.65 | 18.5 |
| TDN | 303.3 | 212.5 | 419.4 | 316 | 301 | 287 | 272 |
| Ash | 108.8 | 89.4 | 68.5 | 78.95 | 84 | 89.03 | 94.1 |
| DE | 5.76 | 3.86 | 8.04 | 5.95 | 5.67 | 5.38 | 5.10 |
| ME | 4.72 | 3.17 | 6.6 | 4.89 | 4.65 | 4.42 | 4.18 |
| GE | 17.17 | 16.79 | 18.7 | 17.8 | 17.6 | 17.4 | 17.2 |
| ADL | 72.3 | 115.7 | 68.9 | 92.3 | 92.7 | 93.2 | 93.6 |

ADL, acid detergent lignin; CHO, carbohydrate; C, commercial; DCP, digestible crude protein; DE, digestible energy; DM, dry matter; EE, ether extract; GE, gross energy; ME, metabolizable energy; NFC, nitrogen free extract; OM, organic matter; TDN, total digestible nutrient; WH, water hyacinth

Conversely, treatment (T4) with the highest inclusion rate of *E. crassipes* exhibited the relatively comparable contents of most of the analyzed nutrients.

### 3.2. Energy and Nutrient intake

The comparative analysis of energy and nutrient intake between sheep and goats fed varying levels of *E. crassipes* is presented in Table 2. Most of the results revealed a distinct variation among species for all nutrient and energy intake parameters (P < 0.001). Sheep consistently exhibited higher intake levels than goats for all nutrients and energy parameters (P <0.05). Among dietary treatment groups for the rams, all nutrient and energy intake parameters were statistically significant (P < 0.001). Specifically, DM, OM, ADL, CHO, GE, and ash contents were significantly lower (P < 0.001) in T1 compared to the other dietary treatment groups. Conversely, DE, DCP, NFC, TDN, ME, and EE contents were significantly lower (P<0.05) in T4 compared to the other dietary treatments groups.

**Table 2.**
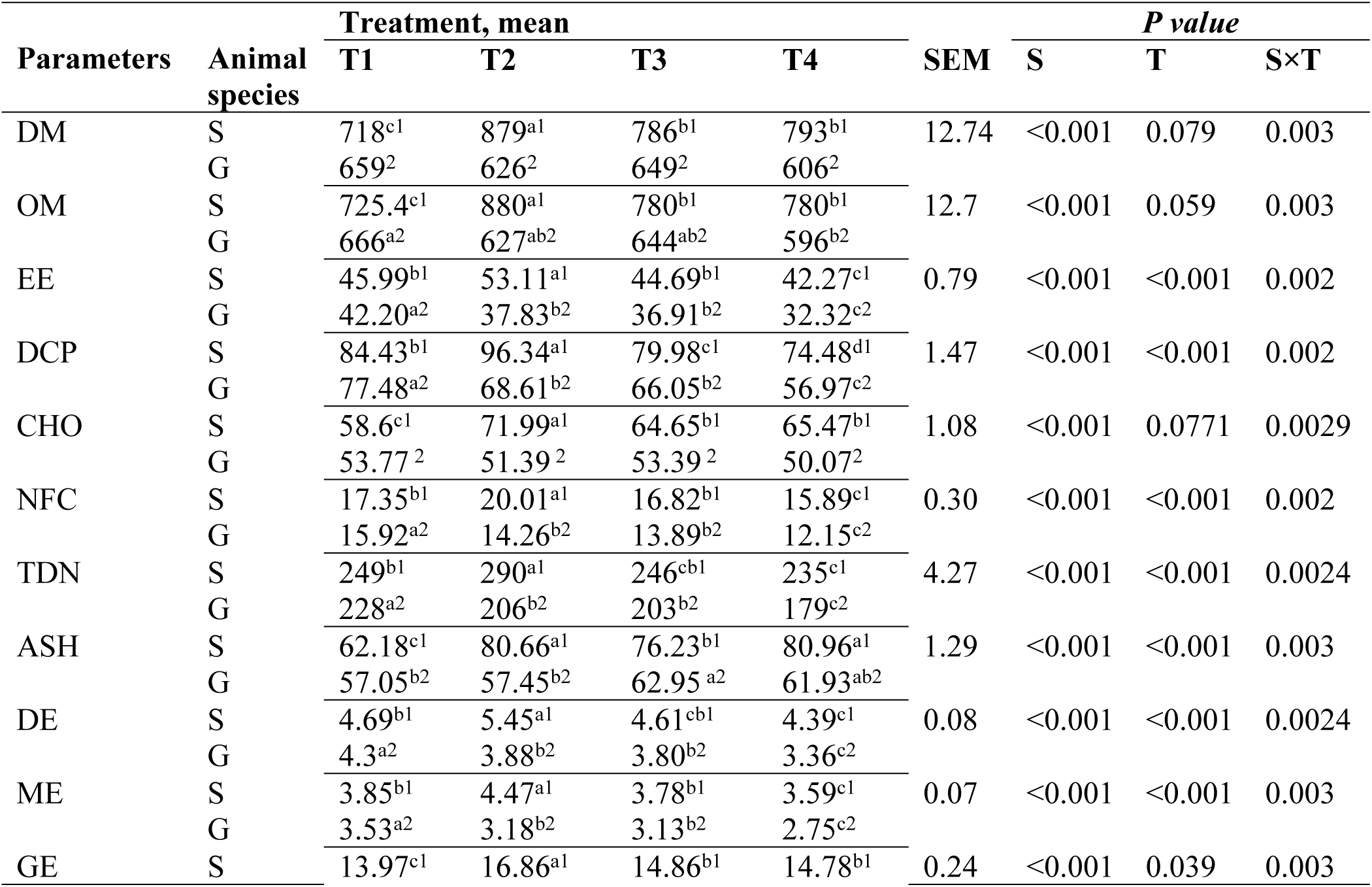
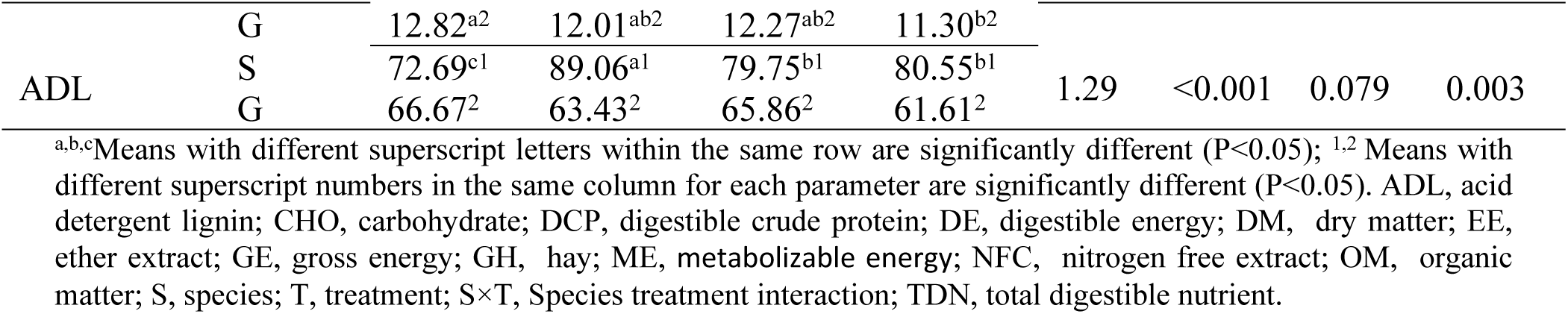
Nutrient and energy intake of sheep and goats.

Additionally, in the dietary treatment groups for the bucks, several nutrient and energy intake parameters were statistically significant (P < 0.001). Notably, EE, DCP, NFC, TDN, DE, ME, and ash content were significantly lower (P < 0.001) in T4 compared to the other treatment groups. On the other hand, OM and GE were significantly lower (P < 0.05) in T4 compared to the other dietary treatments. Furthermore, the result revealed a significant (P < 0.01) interaction effect between species and treatment for all nutrient and energy intake parameters.

### 3.3. Apparent digestibility

The nutrient digestibility coefficients of the experimental diets are presented in Table 3.The results revealed a significant variation in apparent digestibility coefficients of nutrients among the two species. The differences were found to be statistically significant (P < 0.001) for CP, NDF, ADF, and Ash. Likewise, the DM, and OM showed significant variation (P < 0.001) between the two species. Surprisingly, there was significant difference (P < 0.001) across four treatment groups for both species in DM, OM, CP, NDF, ADF, and Ash digestibility. Organic matter (OM) and Dry matter (DM) digestibility was highest in goats, peaking in T1 at 80.16% for DM and 78.49% for OM, and decreasing to 66.84% and 65.13%, respectively, in T4. While sheep exhibited a peak of (75.60% and 73.50%, respectively) T1 but showed a significant decline in later treatments, with the lowest values in T4 (63.67% for DM and 62.36% for OM).

**Table 3.** Apparent digestibility coefficient of nutrients in Doyogena ram and Woyto-Guji buck received different levels of water hyacinth.

| Parameters | Animal Species | Treatment, mean |  |  |  | *P value* |  |  |  |
| --- | --- | --- | --- | --- | --- | --- | --- | --- | --- |
|  |  | T1 | T2 | T3 | T4 | SEM | S | T | S×T |
| OM | S | 75.60<sup>a</sup> | 70.40<sup>b2</sup> | 67.98<sup>b</sup> | 63.67<sup>c2</sup> | 1.16 | <0.001 | <0.001 | 0.638 |
|  | G | 80.16<sup>a</sup> | 76.97<sup>a1</sup> | 72.05<sup>b</sup> | 66.84<sup>c1</sup> |  |  |  |  |
| DM | S | 73.50<sup>a</sup> | 68.80<sup>b1</sup> | 66.40<sup>b</sup> | 62.36<sup>c2</sup> | 1.14 | <0.001 | <0.001 | 0.552 |
|  | G | 78.49<sup>a</sup> | 75.32<sup>a2</sup> | 70.42<sup>b</sup> | 65.13<sup>c1</sup> |  |  |  |  |
| CP | S | 82.60<sup>a2</sup> | 80.69<sup>a</sup> | 73.21<sup>b</sup> | 66.39<sup>c2</sup> | 1.38 | <0.001 | <0.001 | 0.682 |
|  | G | 85.68<sup>a1</sup> | 82.72<sup>b</sup> | 74.89<sup>c</sup> | 69.40<sup>d1</sup> |  |  |  |  |
| NDF | S | 74.53<sup>a2</sup> | 71.42<sup>b2</sup> | 68.29<sup>c2</sup> | 65.42<sup>d</sup> | 0.76 | <0.001 | <0.001 | 0.995 |
|  | G | 76.61<sup>a1</sup> | 73.61<sup>b1</sup> | 70.62<sup>c1</sup> | 67.51<sup>d</sup> |  |  |  |  |
| ADF | S | 60.55<sup>a</sup> | 58.48<sup>b2</sup> | 56.59<sup>c</sup> | 54.22<sup>d</sup> | 0.57 | <0.001 | <0.001 | 0.618 |
|  | G | 62.73<sup>a</sup> | 60.26<sup>b1</sup> | 57.77<sup>c</sup> | 55.41<sup>d</sup> |  |  |  |  |
| ASH | S | 71.45<sup>a2</sup> | 70.70<sup>b2</sup> | 69.48<sup>c2</sup> | 68.62<sup>d2</sup> | 0.32 | <0.001 | <0.001 | 0.673 |
|  | G | 73.43<sup>a1</sup> | 72.73<sup>b1</sup> | 71.67<sup>c1</sup> | 70.37<sup>d1</sup> |  |  |  |  |

<sup>a,b,c</sup> Means with different superscript letters within the same row are significantly different ( $P < 0.05$ ); <sup>1,2</sup> Means with different superscript numbers in the same column for each parameter are significantly different ( $P < 0.05$ ). ADF, acid detergent fiber; CP, digestible crude protein; DM, dry matter; NDF, nutrient detergent fiber, OM, organic matter; S, species; S×T, species treatment interaction; T, treatment

The CP digestibility was highest in goats in T1 (85.68%), while sheep exhibited a peak of 82.60% in T1, with lower digestibility observed in T4 (66.39% for sheep and 69.40% for goats). For NDF and ADF, goats had consistently higher digestibility values, particularly in T1 (76.61% for NDF and 62.73% for ADF, compared to sheep, who showed lower digestibility across all treatments, with values decreasing from 74.53% to 65.42% for NDF and from 60.55% to 54.22% for ADF. Ash digestibility was also higher in goats, peaking at 73.43% in T1, while sheep had lower values, with the highest being 71.45% in T1 and decreasing to 68.62% in T4. It appears that there were no significant (P > 0.05) deviation interactions effect between species and treatment being studied for any of the parameters.

### 3.4. Body weight change and feed conversion efficiency

Table 4 shows the BWC and FCE of the goat bucks and sheep lambs given different dosages of *E. crassipes*. The results revealed a significant variation in body weight change among the two species. The differences were found to be statistically significant (P < 0.001) for IBW and FBW. Likewise, the ADG (g/day) and BWG (kg) showed significant variation (P < 0.05) between the two species. Furthermore, there was significant difference (P < 0.001) across treatment groups for both species in ADG (g/day), BWG (kg) and FCE. Body weight change was highest in sheep, peaking in T1 at 88.09 for ADG (g/day), 1.85 for BWG (kg), 0.11 for FCE and decreasing to 58.73(g/day), 1.23(kg), 0.07, respectively, in T4.

**Table 4.** The trends of body weight change and feed conversion efficiency in Doyogena rams and Woyto-Guji bucks received different levels of *E. crassipes*.

| Parameters | Species | Treatment, Mean |  |  |  | P value |  |  |  |
| --- | --- | --- | --- | --- | --- | --- | --- | --- | --- |
|  |  | T1 | T2 | T3 | T4 | SEM | S | T | S×T |
| IBW (kg) | S | 22.07<sup>1</sup> | 23.07<sup>1</sup> | 22.56<sup>1</sup> | 22.87<sup>1</sup> | 0.32 | <0.001 | 0.884 | 0.938 |
|  | G | 19.12<sup>2</sup> | 19.12<sup>2</sup> | 19.11<sup>2</sup> | 19.94<sup>2</sup> |  |  |  |  |
| FBW(kg) | S | 24.45<sup>1</sup> | 24.68<sup>1</sup> | 23.98<sup>1</sup> | 24.1<sup>1</sup> | 0.31 | <0.001 | 0.059 | 0.554 |
|  | G | 20.70<sup>2</sup> | 20.53<sup>2</sup> | 20.37<sup>2</sup> | 20.70<sup>2</sup> |  |  |  |  |
| ADG (g/day) | S | 88.09<sup>a</sup> | 76.98<sup>ab</sup> | 67.86<sup>ab</sup> | 58.73<sup>b1</sup> | 4.28 | 0.028 | <0.001 | 0.815 |
|  | G | 75.40<sup>a</sup> | 67.46<sup>a</sup> | 59.92<sup>a</sup> | 36.11<sup>b2</sup> |  |  |  |  |
| BWG (kg) | S | 1.85<sup>a</sup> | 1.62<sup>ab</sup> | 1.43<sup>ab</sup> | 1.23<sup>b1</sup> | 0.09 | 0.028 | <0.001 | 0.815 |
|  | G | 1.58<sup>a</sup> | 1.41<sup>a</sup> | 1.26<sup>a</sup> | 0.76<sup>b2</sup> |  |  |  |  |
| FCE | S | 0.11<sup>a</sup> | 0.08<sup>b</sup> | 0.07<sup>b</sup> | 0.07<sup>b</sup> | 0.005 | 0.792 | <0.001 | 0.625 |
|  | G | 0.11<sup>a</sup> | 0.1<sup>a</sup> | 0.08<sup>ab</sup> | 0.06<sup>b</sup> |  |  |  |  |

<sup>a,b,c</sup> Means with different superscript letters within the same row are significantly different (P<0.05); <sup>1,2</sup> Means with different superscript numbers in the same column for each parameter are significantly different (P<0.05). ADG= average daily gain, BWG= body weight gain, FBW= final body weight, FCE= feed conversion efficiency, IBW= initial body weight, S= species, S×T= species treatment interaction, T= treatment

While goat exhibited a peak of (75.40(g/day), 1.58 (kg) and 0.11, respectively) T1 but showed a significant decline in later treatments, with the lowest values in T4 (36.11 for ADG (g/day), 0.76 for BWG (kg) and 0.06 for FCE). On contrary, there were no significant deviation (P > 0.05) interactions effect between species and treatment being studied for any of the parameters.

The body weight change by period in Doyogena sheep and Woyto-guji goat breeds fed varying levels of *E. crassipes* are presented in Table 5. The results revealed a significant variation in body weight change among the two species. The differences were found to be significant (P < 0.001) for IBW, FBW and ADG (g/kg). Likewise, the BWG (kg) showed significant variation (P < 0.05) between the two species. Furthermore, there was significant difference (P < 0.001) across period for both species in FBW, BWG (kg) and FCE. Similarly, the ADG (g/day) showed significant variation (P < 0.05) between the two species.

**Table 5.**
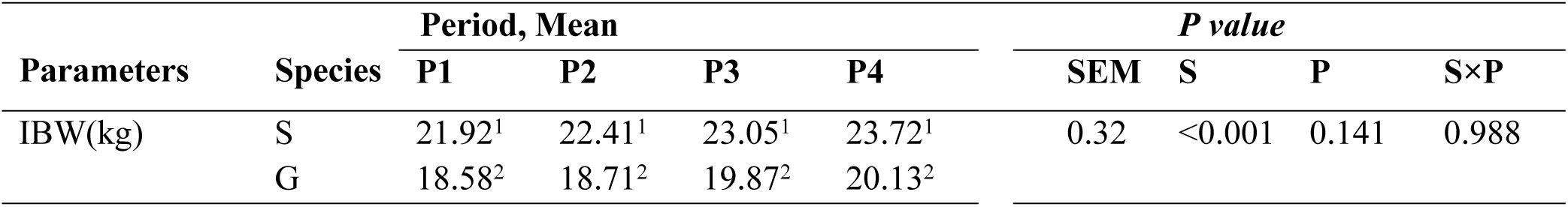
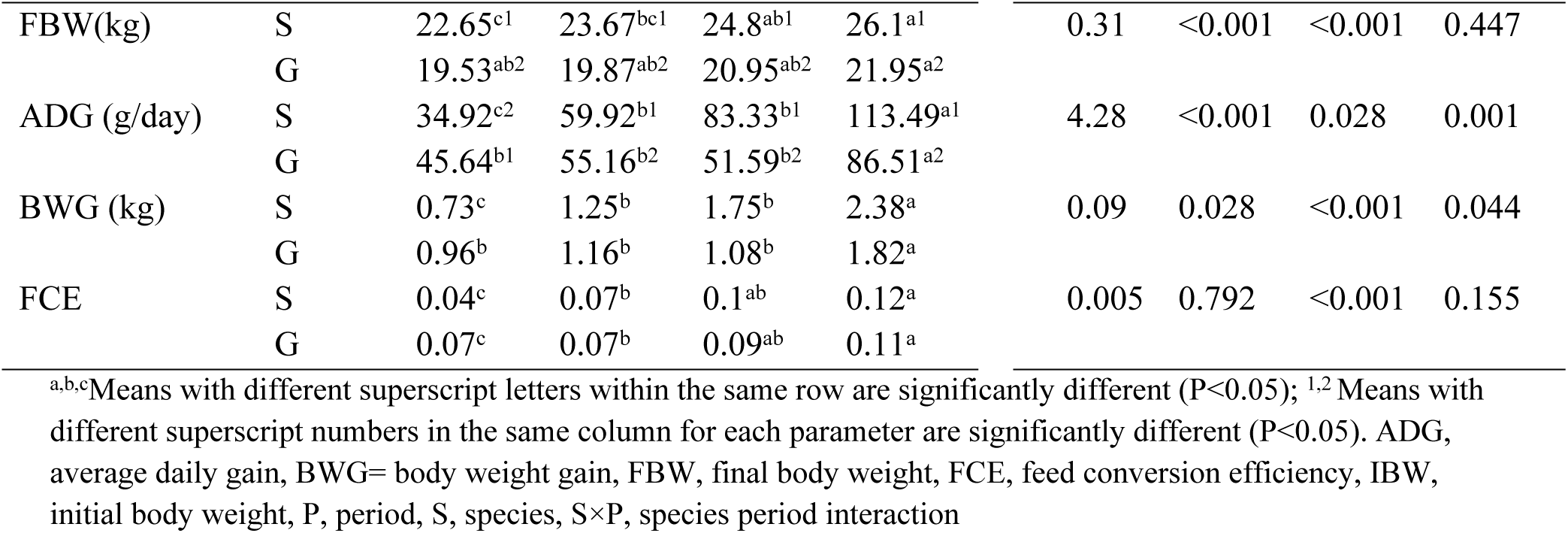
The change in body weight and feed conversion efficiency over different periods in Doyogena rams and Woyto-guji bucks.

| Parameters | Species | Period, Mean |  |  |  | P value |  |  |  |
| --- | --- | --- | --- | --- | --- | --- | --- | --- | --- |
|  |  | P1 | P2 | P3 | P4 | SEM | S | P | S×P |
| IBW(kg) | S | 21.92<sup>1</sup> | 22.41<sup>1</sup> | 23.05<sup>1</sup> | 23.72<sup>1</sup> | 0.32 | <0.001 | 0.141 | 0.988 |
|  | G | 18.58<sup>2</sup> | 18.71<sup>2</sup> | 19.87<sup>2</sup> | 20.13<sup>2</sup> |  |  |  |  |
| FBW(kg) | S | 22.65<sup>c1</sup> | 23.67<sup>bc1</sup> | 24.8<sup>ab1</sup> | 26.1<sup>a1</sup> | 0.31 | <0.001 | <0.001 | 0.447 |
|  | G | 19.53<sup>ab2</sup> | 19.87<sup>ab2</sup> | 20.95<sup>ab2</sup> | 21.95<sup>a2</sup> |  |  |  |  |
| ADG (g/day) | S | 34.92<sup>c2</sup> | 59.92<sup>b1</sup> | 83.33<sup>b1</sup> | 113.49<sup>a1</sup> | 4.28 | <0.001 | 0.028 | 0.001 |
|  | G | 45.64<sup>b1</sup> | 55.16<sup>b2</sup> | 51.59<sup>b2</sup> | 86.51<sup>a2</sup> |  |  |  |  |
| BWG (kg) | S | 0.73<sup>c</sup> | 1.25<sup>b</sup> | 1.75<sup>b</sup> | 2.38<sup>a</sup> | 0.09 | 0.028 | <0.001 | 0.044 |
|  | G | 0.96<sup>b</sup> | 1.16<sup>b</sup> | 1.08<sup>b</sup> | 1.82<sup>a</sup> |  |  |  |  |
| FCE | S | 0.04<sup>c</sup> | 0.07<sup>b</sup> | 0.1<sup>ab</sup> | 0.12<sup>a</sup> | 0.005 | 0.792 | <0.001 | 0.155 |
|  | G | 0.07<sup>c</sup> | 0.07<sup>b</sup> | 0.09<sup>ab</sup> | 0.11<sup>a</sup> |  |  |  |  |

<sup>a,b,c</sup>Means with different superscript letters within the same row are significantly different ( $P < 0.05$ ); <sup>1,2</sup> Means with different superscript numbers in the same column for each parameter are significantly different ( $P < 0.05$ ). ADG, average daily gain, BWG= body weight gain, FBW, final body weight, FCE, feed conversion efficiency, IBW, initial body weight, P, period, S, species, S×P, species period interaction

Body weight change showed the highest in sheep, peaking in P4 at 26.1(kg), 113.49 (g/day), 2.38(kg) and 0.12 for FBW, ADG, BWG and FCE respectively, and decreasing to 22.65(kg), 34.92(g/day), 0.73(kg), and 0.04 for FBW, ADG, BWG and FCE respectively, in P1. While goat exhibited a peak of 21.95(kg), 86.51(g/day), 1.82 (kg) and 0.11 for FBW, ADG, BWG and FCE respectively P4 showed a significant decline in later treatments, with the lowest values in P1 (19.53(kg), 45.64(g/day) 0.73(kg) and 0.07 for FBW, ADG, BWG and FCE respectively. Furthermore, there were significant deviation (P > 0.05) interactions between species and period for ADG (g/day) and BWG (kg).

### 3.5 Economic viability of supplementing water hyacinth in goats and sheep

The Economic viability of supplementing water hyacinth to sheep and goats were presented in table 6. The gross financial margin or total return obtained in this trial was 1,500, 1,600, 1,700 and 1,700 birr/sheep from sheep fed T1, T2, T3 and T4, respectively. 1,200, 1,300, 1,350 and 1,400 birr/goats from goats fed T1, T2, T3 and T4, respectively. The change of net income obtained in T2, T3, T4, and T5 were 41.71, 57.7 and 111.48 birr/ sheep respectively. Whereas, the change of net income obtained in T2, T3, T4, and T5 were 7.17, 14.2 and 21.61 birr/ goats respectively. The marginal rate of return obtained in T2, T3 and T4 were 2, 1.7 and 0.78 birr/ sheep respectively.

**Table 6.** Economic parameters (Birr/ram and Birr/bucks) of Doyo-Gena rams and Woyto-Guji bucks fed diets water hyacinth.

| Parameters | Species | Treatments |  |  |  |
| --- | --- | --- | --- | --- | --- |
|  |  | T1 | T2 | T3 | T4 |
| Initial Price | S | 2500 | 2500 | 2500 | 2500 |
|  | G | 2100 | 2100 | 2050 | 2000 |
| Hay consumed | S | 11.59 | 12.11 | 11.84 | 12.01 |
|  | G | 10.04 | 10.04 | 10.03 | 10.05 |
| Concentrate consumed | S | 11.59 | 9.1 | 5.92 | 3 |
|  | G | 10.04 | 7.53 | 5.015 | 2.51 |
| Water hyacinth consumed | S | 0 | 3.01 | 5.92 | 9.1 |
|  | G | 0 | 2.51 | 5.015 | 7.54 |
| **Cost** |  |  |  |  |  |
| Concentrate (ETB/sheep) | S | 278.16 | 218.4 | 142.08 | 72 |
|  | G | 240.96 | 180.72 | 120.36 | 60.24 |
| Grass hay (ETB/sheep) | S | 124.18 | 129.75 | 126.86 | 128.67 |
|  | G | 107.57 | 107.57 | 107.57 | 107.57 |
| Water hyacinth (ETB/sheep) | S | 0 | 20.9 | 41.1 | 63.15 |
|  | G | 0 | 17.41 | 34.80 | 52.33 |
| Labor (ETB/sheep) | S | 62.5 | 62.5 | 62.5 | 62.5 |
|  | G | 62.5 | 62.5 | 62.5 | 62.5 |
| Medication (ETB/sheep) | S | 50 | 50 | 50 | 50 |
|  | G | 50 | 50 | 50 | 50 |
| Selling price (ETB/sheep) | S | 4,000 | 4,100 | 4,200 | 4,200 |
|  | G | 3,300 | 3,400 | 3,400 | 3,400 |
| TR(A) | S | 1,500 | 1,600 | 1,700 | 1,700 |
|  | G | 1,200 | 1,300 | 1,350 | 1,400 |
| $\Delta TR$ | S | 0 | 100 | 200 | 200 |
|  | G | 0 | 100 | 150 | 200 |
| TVC (B) | S | 514.84 | 481.55 | 422.54 | 376.32 |
|  | G | 461.03 | 418.2 | 375.23 | 332.64 |
| $\Delta TVC$ | S | 0 | 33.29 | 92.3 | 138.52 |
|  | G | 0 | 42.83 | 85.8 | 128.39 |
| NI (A-B) | S | 985.16 | 1,118.45 | 1,277.46 | 1,323.68 |
|  | G | 738.97 | 881.8 | 974.77 | 1,067.36 |
| $\Delta NI (\Delta TR - \Delta TVC)$ | S | 0 | 66.71 | 107.7 | 107.7 |
|  | G | 0 | 57.17 | 64.2 | 71.61 |
| $MRR = \Delta NI / \Delta TVC$ | S | 0 | 2 | 1.17 | 0.78 |
|  | G | 0 | 1.34 | 0.75 | 0.56 |

S= sheep; G= goats; NI= net income; TR= total return; TVC=total variable cost; MRR= marginal rate of return; ETB= Ethiopian currency (Birr)

## 4. DISCUSSION

### 4.1. Nutritional values of the experimental diet

In our study, the energy content of the experimental diet, encompassing DCP, GE, TDN, ME, DE and total CHO, generally aligned within the expected range as outlined in existing literature. Nevertheless, some discrepancies with findings from other studies were observed. Specifically, our DCP values were higher than those reported by Tavirimirwa *et al*. (2012) at 31.9 g/kg DM and 29.6 g/kg DM, but lower than the 71.76 g/kg DM reported by Gashaw and Defar (2017) and closest to the 51.9 g/kg DM and 49.5 g/kg DM reported by Gobena *et al*. (2022). The GE content in our study was comparable to the 17.29 MJ/kg DM reported by Quintero-Anzueta *et al*. (2021), yet lower than the 18.96 –19.06 MJ/kg DM range found by Pathot and Berhanu (2023) and higher than the 15.52 MJ/kg DM reported by Mekuriaw *et al*. (2020).

The TDN content in our study exceeded the 102.2 g/kg DM reported by Fekadu *et al*. (2018) but was less than the 377 g/kg DM reported by Singh *et al*. (2024). Our DE and ME values were similar to the 4.72 MJ/kg DM and 3.87 MJ/kg DM reported by Chino Velasquez *et al*. (2022), although they were notably lower than the ranges of DE 6.8-8.9 MJ/kg DM and ME 5.6-7.3 MJ/kg DM reported by Carvajal-Tapia *et al*. (2023). The TCHO content aligned with the 80.8 g/kg DM reported by Singh *et al*. (2024) but was higher than the 64.38-67.91 g/kg DM range reported by Pathot and Berhanu (2023). This variation in energy content may be due to factors such as the grass species, environmental conditions like soil fertility and climate, as well as aspects such as maturity, growing conditions, post-harvest practices, and overall field management.

For water hyacinth, the DCP value observed in this study was in line 104.1 g/kg DM and 113.1 g/kg DM with findings from Hossain *et al*. (2015) and Akinwande *et al*. (2013) respectively. It was higher than the values of 93.3, 93.7, and 73.3 g/kg DM reported by Mako *et al*. (2011, 2012) and Suleiman *et al*. (2020), then lower than the findings of 196.4 and 130.9 reported by Enyi *et al*. (2020) and Mekuriaw *et al*. (2018), respectively. The total CHO value in our study was in accord with the 71.4 reported by Suleiman *et al*. (2020), higher than the 68.6 g/kg DM, 63.3 g/kg DM, 63.12 g/kg DM, and 67.23 g/kg DM reported by Mako *et al*. (2011, 2012), Akinwande *et al*. (2013), and Enyi *et al*. (2020), however lower than the 74.6 g/kg DM reported by Hossain et al. (2015).

The TDN value found in this study is to some extent closer to the value 276.45 g/kg DM reported by Enyi *et al*. in 2020. However, it surpasses the values of 189.65 g/kg DM, 194.2 g/kg DM, 193.87 g/kg DM, and 219.4 g/kg DM reported by Suleiman et al. (2020), Mako *et al*. (2011 and 2012), and Hossain *et al*. (2015). In contrast, this result is significantly lower than the higher values of 407.7 g/kg DM and 461 g/kg DM reported by Akinwande *et al*. (2013) and de Vasconcelos *et al*. (2016). The DE value obtained in this study in line with the 6.14 MJ/kg DM conveyed by Enyi *et al*. (2020). However, this value is pointedly higher than the 3.7 MJ/kg DM reported by Suleiman *et al*. (2020). Conversely, it is notably lower than the 10.2 MJ/kg DM reported by Hossain *et al*. (2015). The ME value for water hyacinth in this study aligns most closely with the 5.03 MJ/kg DM reported by Enyi *et al*. (2020). However, this result is higher than the 3.03 MJ/kg DM reported by Suleiman et al. (2020). Conversely, our ME value is lower than the 8.37 MJ/kg DM reported by Hossain *et al*. (2015). The GE value obtained in this study for water hyacinth is similar to the result 17.4 MJ/kg DM reported by Akinwande *et al*. (2013). However, this result is higher compared to that of 14.9 MJ/kg DM as reported by Mako *et al*. (2012). This difference in energy content probably attributed to factors such as aquatic habitat conditions, seasonal variations, and frequency of harvesting.

### 4.2. Energy and Nutrient intake of goats and sheep

The overall nutrient and energy intake of sheep and goats varied pointedly across treatments. The observed higher nutrient and energy intake in sheep compared to goats possibly attributed to their distinct feeding habits. Sheep, being grazers, generally consume a wider variety of forage and are less selective about their diet, leading to higher intake levels. Conversely, goats, known as browsers, tend to be more selective in their feeding, particularly in confined environments, resulting in lower overall nutrient and energy consumption. It is encouraging to compare this figure with that found by scholars (Salim *et al*., 2002; Bahtti, 2013 and Candyrine *et al*., 2019) they find that higher nutrient intake by sheep compared to goats. In addition to this notably, T2 in sheep and T1 in goats exhibit the highest in all nutrient and energy parameters mean value, significantly outperforming the other treatments. On the flip side, as the inclusion rate of water hyacinth increases, the energy values such as DCP, TDN, DE, ME, and GE show a consistent numerical decline among T1, T2, T3, and T4. Our results align with de Vasconcelos *et al*. (2016), who found that replacing Tifton-85 hay with WH hay resulted in a linear decrease in DM, OM, CP, and NDF intake, with the most significant decline observed after certain level of higher water hyacinth hay inclusion. Similarly, Mekuriaw *et al*. (2018) mirror for our findings, who reported significant decline in DM, OM, CP and NDF intake while increasing water hyacinth hay after reaching a certain level of replacement.

### 4.3. Apparent digestibility

Among all treatment groups, Woyto-Guji goats show significant difference in digestibility for all parameters like DM, OM, CP, NDF, ADF, and Ash, suggesting a more efficient utilization of these nutrients compared to doyogena sheep. Lamy *et al*. (2011) reported that goats are higher digestion coefficients compared with sheep across various treatment groups. Similarly, Kechero *et al*., 2015 and Kijas *et al*. (2012), who reported goats, are more efficient in the digestion of fiber and the utilization of poor roughages than sheep. These variations are likely, as a result of the distinct digestive and physiological mechanisms like rumen function, digestive enzyme activity, or microbial flora that each species uses to process and absorb nutrients from their diet. Conversely, as the inclusion rate of water hyacinth rises, the coefficient of all studied nutrient digestion regardless of the species exhibit significantly decline among T1, T2, T3, and T4. This study was in line with the result reported by Mekuriaw *et al*. (2018), as the level water hyacinth leaves inclusion rate rise DM, OM, CP and NDF coefficient of digestion decrease among T1, T2, T3, and T4. Similarly, Mengistu *et al*. (2020), reported as the level of Dried Mulberry and *Vernonia amygdalina* mixed leaves inclusion rate rise in a diet, a significant decline in coefficient of digestion for DM, OM, CP, NDF and ADF among T1, T2, T3, and T4. This was probably, attributed to the lower CP and higher ADF and NDF contents of the water hyacinth compared to commercial concentrate.

### 4.4. Body weight change and feed conversion efficiency

Noticeably, IBW was significantly different between Doyogena sheep and Woyto-Guji goats, before commencing the experiment, as expected, there were significant differences in FBW, ADG, and BWG in the current study. Our finding corroborates with Candyrine *et al*. (2019) who reported higher FBW, ADG and BWG in sheep compared to goats. This difference is likely species-specific due to grazers tending to consume a wider range of available feed. In contrast, browsers are more selective about their feed choices. Similarly, confinement pen feeding used in the study may also favor these feeding behaviors, leading to better body weight change in sheep. The FBW of the Doyogena sheep and Woyto-Guji goats increased during the feeding period for all treatments in a way that was consistent with their intakes of total dry matter. The findings of current study agrees with the report of Mekuriaw *et al*. (2018), who found that supplementation with water hyacinth leaf meal as concentrate replacer had no significant effect on the final weight of Washera sheep of Ethiopian highlands.

The lowest ADG, BWG, and FCE recorded in Doyogena sheep and Woyto-Guji goats supplemented T4 which was 35% water hyacinth inclusion. Similarly, higher ADG, BWG and FCE obtained in this study were in T2 and T3 than T4. This was probably attributed to the low intake of water hyacinth, which may be allied to the high nutrient detergent fiber content of the plant. The highest ADG, BWG, and FCE recorded in Doyogena sheep and Woyto-Guji goats supplemented T1 concentrate and basal diet alone. The discrepancy regardless of the species in ADG, BWG, and FCE between dietary treatment groups could be attributed to differences in intake of the diet and nutritive value in the diets. This study was in accord with the result stated by Khalid *et al*. (2012) quality and quantity of nutrients in the feed directly influence the efficiency and productivity of ruminant animals.

The ADG of sheep in this study were comparable with that reported at 62 g/day to 87 g/day for Arsi-Bale sheep in Ethiopia fed a grass hay basal diet supplemented with malted oat grain and noug seed cake (Girma *et al*., 2014). However, lower than 138 g/day –189 g/day for small-tail Han lambs fed with different ratios of alfalfa hay and maize stover (Sun *et al*., 2018). Whereas, the ADG of goats in current study were comparable with that reported at 76.2 g/day–59.71 g/day for Borana goats fed a grass-hay basal diet supplemented with three browse species mixed with wheat bran (Kumsa *et al*., 2019). Though higher than 30 g/day for local goats fed urea-treated *tef* straw basal diet supplemented cassava leaf meal, brewers’ dried grain, and their mixture (Tilahun *et al*., 2013).

### 4.5 Economic viability of supplementing water hyacinth in goats and sheep

The partial replacement 35% WH + 15% CC (T4), followed by 25% WH + 25% CC (T3) and 15% WH + 35% CC (T2) with mixed grass hay for Woyito-Guji bucks and Doyo-Gena rams had a positive effect in reducing the production cost. The group fed on diet with a higher level of 35% WH + 15% CC (T4) achieved the highest positive net income and profit margin 0% WH + 50% CC ( T1). Compared to the control diet, feed cost was reduced by 37.2% (420.34 vs 263.82 Birr/ram) and 36.8% (348.53 vs 220.14 Birr/buck) in the group fed on 35% WH + 15% CC (T4). In comparison to T1, the feed cost share of the ram in TVC was decreased by roughly 1.5, 4.7, and 8% in T2, T3, and T4, respectively. Similarly, the feed cost share of the TVC ram was reduced in T2, T3, and T4 by approximately 2.5, 5.6, and 9.4%, respectively, when compared to T1. Both species had higher net income in T4 and increased amounts of water hyacinth meal substituted for concentrate mixture because it is less expensive than concentrate feeds. The cost per kilogram of feed decreased as the amount of water hyacinth meal in the diets rose. The treatment with the highest net return above control has the highest potential for financial gain. Greater profitability or economic benefit with an increase in the water hyacinth meal replacement level on concentrate feed is shown by increases in the marginal return index. In this particular research, 35% water hyacinth meal replacement in T4 improved body weight change of sheep and goat correspondingly increased net income from the sale of sheep at the end of experimental period. Thus, at the anticipated cost found in this study, T4 would be suggested as the biological and economical feeding regimen for developing marketable Woyito-Guji bucks and Doyogena rams.

## 5. CONCLUSION

The studied parameters disclosed significant variation across species in nutrient utilization efficiency, particularly nutrient and energy intake, nutrient digestibility, average daily gain, and body weight gain. Notably, sheep exhibited the highest values in terms of feed intake, while goats had the lowest among the treatment groups consuming similar diets. There were also significant variations in nutrient digestibility, ADG, BWG, and FCE among treatment groups for both species. Moreover, for findings of energy and nutrient digestibility, similar trends were exhibited for nutrient and energy intake among treatment groups for both species, with few exceptions, particularly in goats. Furthermore, there were significant interactions between species and treatment across all nutrient and energy intake parameters. Based on a comprehensive analysis, feeding regimen T4 is recommended as the optimal dietary approach for maximizing both biological growth and economic efficiency in raising Woyito-Guji bucks and Doyogena rams. The current study concludes that supplementing Woyito-Guji bucks and Doyogena rams with water hyacinth—replacing up to 37.5% of the commercial concentrate mix—can enhance nutrient intake, digestibility, and body weight gain. This aquatic weed has the potential to serve as an alternative protein supplement in a natural pasture hay-based feeding system for these breeds during the dry season. Substantial quantities of water hyacinth are available in the study area, when traditional forages often fall short in providing sufficient nitrogen for optimal rumen function in ruminants.

### Ethics approval

The Animal Research Ethics Committee of Arba Minch University approved the research methodologies and procedures, verifying that they follow ethical standards. The committee issued a certificate with ref. no AMU/AREC/4/2016 stating that the study was approved and carried out in an ethical and responsible manner.

## Acknowledgements

The authors would like to express their appreciation to the Arba Minch University VLIR-IUC project for funding their research work. They would also like to acknowledge the Arba Minch University Animal Research Farm and Chemistry Laboratory for their technical support in analyzing the chemical composition of the feed samples, respectively. Furthermore, the authors would like to extend their gratitude to all those who have contributed to the realization of this article.

## Conflict of interest

The authors declare that there are no conflicts of interest.

## Notes

### Competing Interest Statement

The authors have declared no competing interest.

